# Haplotype-Based Noninvasive Prenatal Diagnosis for Duchenne Muscular Dystrophy: A pilot study in South China

**DOI:** 10.1101/551200

**Authors:** Min Chen, Chao Chen, Yingting Li, Yuan Yuan, Zhengfei Lai, Fengyu Guo, Yaoshen Wang, Xiaoyan Huang, Shiquan Li, Renhua Wu, Zhiyu Peng, Jun Sun, Dunjin Chen

## Abstract

**Objective:** To explore the accuracy and feasibility of noninvasive prenatal diagnosis (NIPD) for Duchenne Muscular Dystrophy (DMD) based on the haplotype approach.

**Methods:** We recruited singleton pregnancies at-risk of DMD at 12-25 weeks of gestation from 17 families who all had a proband children affected by DMD. We have identified the pathogenic mutations in probands and their mothers by multiplex ligation-dependent probe amplification (MLPA). To construct parental haplotypes, we performed captured sequencing on genomic DNA from parents and probands. The integration analysis of parental haplotypes and targeted sequencing results of maternal plasma DNA were used to infer the fetal haplotype and genotypes in *DMD* gene. Fetal *DMD* genotypes were further confirmed by invasive prenatal diagnosis.

**Results:** We have successfully performed the haplotype-based NIPD in all recruited families. Ten fetuses were identified as normal, including four female and six male fetuses. Four female fetuses were carriers and the other three male fetuses were affected by *DMD* with exons 49-52 deletion, exons 8-37 deletion and c.628G > T mutation, respectively. The results of NIPD were consistent with those of invasive diagnosis.

**Conclusion:** Haplotype-based NIPD for DMD by targeted sequencing is promising and has potential for clinical application.

## Introduction

Duchenne Muscular Dystrophy (DMD) is an X-linked recessive genetic disorder affecting 1 in 3,500 new-born males. Approximately 70% DMD patients have at least one exon deletions/repeats, and the others are point mutations or small insertions/deletions of *DMD* gene. The current standard in prenatal diagnosis is to provide invasive procedures, chorionic villus sampling (CVS) or amniocentesis for genetic study[1]. However, a small risk of miscarriage is associated with invasive procedures[2].

During the last decade, non-invasive prenatal test (NIPT) for aneuploidy using maternal plasma cell free fetal DNA (cff-DNA) has been widely used in clinical practice. Further research has been conducted to develop NIPD for single gene disorders (SGDs) using various technologies such as real-time PCR, COLD-PCR, digital PCR, cSMART and next-generation sequencing (NGS).The initial attempts were limited to the exclusion of paternal inheritance[3, 4] and detection of de novo mutations[5]. The relative haplotype approach has been shown to be a perfect solution to detect the maternal inherited alleles and alleles shared by both parents[6-8]. However, establishing an accredited NIPD service is challenging with regards to the costs-effectiveness and quality control of fetal DNA fraction, reads depth of cff-DNA and the informative SNPs[9].

Here we reported the haplotype-based NIPD for DMD in a large sample size.

## Material and methods

### Patients and sample collection

Seventeen at-risk families with probands (male) were recruited with genetic counseling and informed consent. The study was approved by the institutional Ethics Committee. For each family, we collected 10ml blood samples from the parents and proband, 5mg chorionic villus (CV) or 10ml amniotic fluid (AF) from intrauterine cavity. The causative mutation in *DMD* gene in each family was identified using MLPA (Table 1).

**Table 1.** Clinical Information and Molecular Diagnosis

| Family | Sample | Clinical | Age | GW | DMD MLPADiagnosis | PregancyOutcome |
| --- | --- | --- | --- | --- | --- | --- |
| F01 | mother |  |  |  | Het. D.EX46-48 |  |
|  | father |  |  |  | Normal |  |
|  | proband |  |  |  | Male, D.EX46-48 |  |
|  | plasma |  |  | 13W+1D | □ |  |
|  | CV |  |  |  | Female, Het. D.EX46-48 |  |
| F02 | mother |  |  |  | Het. D.EX20-37 |  |
|  | father |  |  |  | Normal |  |
|  | proband |  |  |  | Male, D.EX20-37 |  |
|  | plasma |  |  | 18W+1D | □ |  |
|  | AF |  |  |  | Female, Normal |  |
| F03 | mother |  |  |  | Het. D.EX7-47 |  |
|  | father |  |  |  | Normal |  |
|  | proband |  |  |  | Male, D.EX7-47 |  |
|  | plasma |  |  | 11W+3D | □ |  |
|  | CV |  |  |  | Male, Normal |  |
| F04 | mother |  |  |  | Het. D.EX43-45 |  |
|  | father |  |  |  | Normal |  |
|  | proband |  |  |  | Male, D.EX43-45 |  |
|  | plasma |  |  | 20W+1D | □ |  |
|  | AF |  |  |  | Female, Het. D.EX43-45 |  |
| F05 | mother |  |  |  | Het. D.EX45-47 |  |
|  | father |  |  |  | Normal |  |
|  | proband |  |  |  | Male, D.EX45-47 |  |
|  | plasma |  |  | 14W+1D | □ |  |
|  | CV |  |  |  | Female, Normal |  |
| F06 | mother |  |  |  | Het. D.EX49-52 |  |
|  | father |  |  |  | Normal |  |
|  | proband |  |  |  | Male, D.EX49-52 |  |
|  | plasma |  |  |  | □ |  |
|  | CV |  |  | 12W+1D | Male, D.EX49-52 |  |
| F07 | mother |  |  |  | Het.c.628G>T |  |
|  | father |  |  |  | Normal |  |
|  | proband |  |  |  | Male, c.628G>T |  |
|  | plasma |  |  | 13W+2D | □ |  |
|  | CV |  |  |  | Male, c.628G>T |  |
| F08 | mother |  |  |  | Het. D.EX17 |  |
|  | father |  |  |  | Normal |  |
|  | proband |  |  |  | Male, D.EX17 |  |
|  | plasma |  |  | 13W+2D | □ |  |
|  | CV |  |  |  | Female, Normal |  |
| F09 | mother |  |  |  | Het. D.EX45-48 |  |
|  | father |  |  |  | Normal |  |
|  | proband |  |  |  | Male, D.EX45-48 |  |
|  | plasma |  |  | 19W+1D | □ |  |
|  | AF |  |  |  | Female, Het. D.EX45-48 |  |
| F10 | mother |  |  |  | Het. D.EX8-37 |  |
|  | father |  |  |  | Normal |  |
|  | proband |  |  |  | Male, D.EX8-37 |  |
|  | plasma |  |  | 12W+1D | ☐ |  |
|  | CV |  |  |  | Male, D.EX8-37 |  |
| F11 | mother |  |  |  | Het. D.EX54 |  |
|  | father |  |  |  | Normal |  |
|  | proband |  |  |  | Male, D.EX54 |  |
|  | plasma |  |  | 12W+5D | ☐ |  |
|  | CV |  |  |  | Male, Normal |  |
| F12 | mother |  |  |  | Het. D.EX48-49 |  |
|  | father |  |  |  | Normal |  |
|  | proband |  |  |  | Male, D.EX48-49 |  |
|  | plasma |  |  | 11W+2D | ☐ |  |
|  | CV |  |  |  | Female, Normal |  |
| F13 | mother |  |  |  | Het. Dup.EX49-50 |  |
|  | father |  |  |  | Normal |  |
|  | proband |  |  |  | Male, Dup.EX49-50 |  |
|  | plasma |  |  | 12W+4D | ☐ |  |
|  | CV |  |  |  | Male, Normal |  |
| F14 | mother |  |  |  | Het. D.EX46-50 |  |
|  | father |  |  |  | Normal |  |
|  | proband |  |  |  | Male, D.EX46-50 |  |
|  | plasma |  |  | 26W+6D | ☐ |  |
|  | AF |  |  |  | Male, Normal |  |
| F15 | mother |  |  |  | Het. D.EX12-20 |  |
|  | father |  |  |  | Normal |  |
|  | proband |  |  |  | Male, D.EX12-20 |  |
|  | plasma |  |  | 20W+3D | ☐ |  |
|  | AF |  |  |  | Male, Normal |  |
| F16 | mother |  |  |  | Het.c.1408A>T |  |
|  | father |  |  |  | Normal |  |
|  | proband |  |  |  | Male, c.1408A>T |  |
|  | plasma |  |  | 25W+4D | ☐ |  |
|  | AF |  |  |  | Male, Normal |  |
| F17 | mother |  |  |  | Het. D.EX14-15 |  |
|  | father |  |  |  | Normal |  |
|  | proband |  |  |  | Male, D.EX14-15 |  |
|  | plasma |  |  | 11W+1D | ☐ |  |
|  | CV |  |  |  | Female, Het. D.EX14-15 |  |

Notes: Abbreviations: CV, Chorionic Villi ;AF, Amniotic fluid; Het. D.EX, Heterozygous deletion Exon; Y, Years; GW, Gestational Weeks.

### DNA sequencing library preparation

A 657.29Kb SeqCap kit (Roche, Basel, Switzerland) containing 13.91Kb coding region, 3965 single nucleotide polymorphisms (SNPs) (MAF 0.3-0.5) located within the 1M region flanking *DMD* gene (Figure 1) and gender determination locus in chromosome Y was designed. Genomic DNA (gDNA) was extracted from blood and AF or CV sample with QIAamp DNA Blood Mini Kit. Cff-DNA was extracted from plasma performing QIAamp Circulating Nucleic Acid kit. Cff-DNA and gDNA library was prepared referred to Kapa Biosysterm library preparation kit and Illumina standard protocol, respectively. The post-capture libraries were sequenced by PE 101 bp on illumina Hiseq2500.

**Figure 1.**
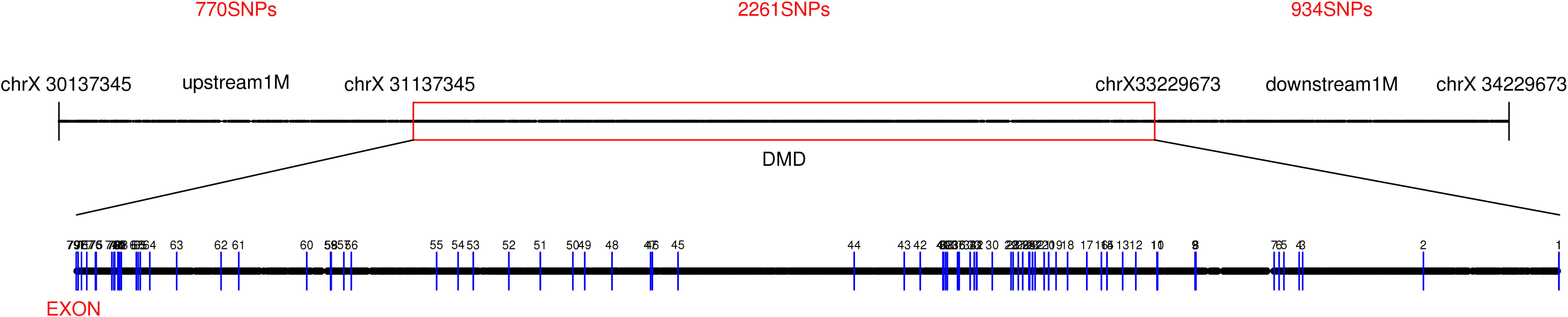
Target region of *DMD* gene and SNPs used for haplotyping.

### Haplotype-based NIPD for DMD

After sequencing, the analysis of raw data, calculation of fetal DNA fraction and plasma sequence error ratio could be processed according to the bioinformatics pipeline reported before [10]. The average depth and coverage of the specific region on Y chromosome was used to infer fetal gender. Plasma samples of male fetus demonstrated 4 times higher coverage on target region of Y chromosome and higher mean depth but almost no reads mapped to Y specific region in female fetus. Thus, the sex of the fetus could be determined based on the average depth and coverage of the specific region of Y chromosome.

The haplotype linked with mutant and wild allele was constructed using the sequence data of family (Figure 2). Based on the linkage relationship obtained from parental haplotypes and the base sequence obtained from plasma DNA sequencing, the Hidden Markov Model (HMM) was constructed to deduce the fetal genetic allele of *DMD*. Finally, we used the Viterbi algorithm to infer the most likely inherited haplotype. To construct the HMM, we analyzed the SNPs where mother was heterozygous and father was homozygous. For each site, we used the number of reads in maternal plasma to calculate the probabilities that the fetus inherited the pathogenic and non-pathogenic allele. These probabilities were HMM emission probabilities. Recombination rates between SNPs are provided via a genetic map (from NCBI) that specifies the genetic position of the SNPs in cM, and these probabilities were HMM transition probabilities.

**Figure 2.**
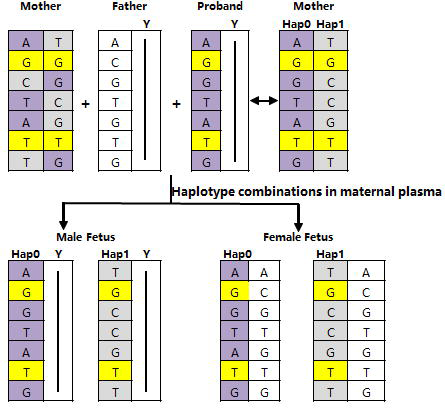
The haplotype-based approach of NIPD for DMD.

### Validation of NIPD for DMD

The *DMD* genotype of fetal sample obtained by invasive procedure was confirmed by MLPA (SALSA P021 and P060 DMD probe mix, Netherlands) in a blind manner. The AF or CV sample was also captured sequencing using the same probe to further prove the accuracy of NIPD. To evaluate the accuracy of NIPD under different sequencing depth/fraction/informative of SNPs, a series of computer depth/fraction simulation experiment was executed by comparing the inferred SNPs with the fetal genotyping.

## Results

The clinical information of seventeen families including the fetal genotype was shown in Table 1. *DMD* mutations in each family including large fragment deletions and point variants are summarized in Table 1.

After sequencing and bioinformatics analysises, a mean of 152.51x and 248.80x reads was obtained for gDNA and cell free DNA, respectively (Table 2). The mean (± SD) capture specificity, 20xcoverage and duplication rate in the target region were 51.60% (±8.62%), 98.83% (±0.92%) and 29.39% (±6.09%), respectively (Table 2).

**Table 2.** Statistics of Target Region Sequencing Data

| Case | Sample ID | Capture specificity | Duplication rate | Depth of target region | Coverage (Depth >= 20X) | Mean depth of target region in ChrY | Coverage $\geq 4\times$ in ChrY | Type 1 SNPs (n) | Type 2 SNPs (n) | Error rate | cffDNA concentration |
| --- | --- | --- | --- | --- | --- | --- | --- | --- | --- | --- | --- |
| F01 | mother | 42.38% | 16.80% | 159.79 | 99.08% | 1.50 | 26.70% |  |  |  |  |
|  | father | 60.28% | 17.93% | 149.98 | 98.82% | 105.35 | 100.00% |  |  |  |  |
|  | proband | 42.05% | 18.08% | 159.00 | 98.75% | 112.37 | 100.00% |  |  |  |  |
|  | plasma | 41.37% | 19.63% | 158.92 | 99.13% | 5.87 | 29.21% | 484 | 470 | 0.35% | 9.30% |
|  | CV | 74.16% | 45.15% | 479.31 | 99.63% | 1.23 | 25.84% |  |  |  |  |
| F02 | mother | 47.87% | 27.98% | 116.24 | 98.55% | 0.00 | 0.00% |  |  |  |  |
|  | father | 52.97% | 32.29% | 125.85 | 98.66% | 81.75 | 100.00% |  |  |  |  |
|  | proband | 53.43% | 27.69% | 106.24 | 97.73% | 68.85 | 100.00% |  |  |  |  |
|  | plasma | 53.97% | 29.26% | 116.45 | 98.52% | 2.49 | 10.05% | 329 | 592 | 0.33% | 8.50% |
|  | AF | 58.08% | 33.50% | 213.47 | 99.45% | 1.57 | 22.84% |  |  |  |  |
| F03 | mother | 69.33% | 27.54% | 118.81 | 98.59% | 0.00 | 0.00% |  |  |  |  |
|  | father | 57.65% | 28.44% | 97.86 | 97.81% | 64.21 | 100.00% |  |  |  |  |
|  | proband | 54.54% | 26.92% | 94.11 | 94.37% | 63.21 | 100.00% |  |  |  |  |
|  | plasma | 49.07% | 32.89% | 112.57 | 98.57% | 26.33 | 99.28% | 402 | 221 | 0.28% | 12.77% |
|  | CV | 53.45% | 32.10% | 262.80 | 99.67% | 65.46 | 100.00% |  |  |  |  |
| F04 | mother | 51.56% | 19.04% | 78.25 | 97.24% | 1.13 | 32.39% |  |  |  |  |
|  | father | 50.35% | 27.03% | 137.38 | 98.86% | 91.57 | 100.00% |  |  |  |  |
|  | proband | 49.58% | 24.59% | 86.80 | 96.45% | 56.51 | 100.00% |  |  |  |  |
|  | plasma | 53.10% | 28.46% | 223.09 | 99.16% | 2.63 | 9.89% | 600 | 410 | 0.38% | 7.83% |
|  | AF | 56.59% | 39.03% | 208.07 | 99.44% | 1.68 | 41.18% |  |  |  |  |
| F05 | mother | 53.10% | 28.34% | 149.61 | 99.05% | 0.00 | 0.00% |  |  |  |  |
|  | father | 54.90% | 26.45% | 83.57 | 97.17% | 56.20 | 99.86% |  |  |  |  |
|  | proband | 49.23% | 27.00% | 78.85 | 96.33% | 52.37 | 100.00% |  |  |  |  |
|  | plasma | 52.51% | 30.98% | 78.09 | 97.00% | 2.06 | 38.70% | 460 | 312 | 1.92% | 11.15% |
|  | CV | 74.09% | 19.40% | 264.05 | 99.45% | 1.10 | 16.03% |  |  |  |  |
| F06 | mother | 57.28% | 28.35% | 113.72 | 98.55% | 0.00 | 0.00% |  |  |  |  |
|  | father | 54.44% | 28.35% | 98.23 | 97.98% | 66.87 | 99.98% |  |  |  |  |
|  | proband | 69.57% | 33.59% | 106.73 | 97.72% | 73.24 | 100.00% |  |  |  |  |
|  | plasma | 55.26% | 32.32% | 90.35 | 97.85% | 14.11 | 99.13% | 317 | 269 | 2.16% | 6.22% |
|  | CV | 46.40% | 22.56% | 275.98 | 99.48% | 32.29 | 100.00% |  |  |  |  |
| F07 | mother | 57.04% | 29.49% | 108.09 | 98.36% | 0.00 | 0.00% |  |  |  |  |
|  | father | 56.63% | 32.15% | 128.55 | 98.81% | 84.56 | 100.00% |  |  |  |  |
|  | proband | 55.05% | 30.52% | 106.56 | 98.17% | 68.63 | 100.00% |  |  |  |  |
|  | plasma | 53.58% | 34.84% | 148.73 | 99.05% | 18.45 | 99.39% | 419 | 450 | 0.39% | 9.42% |
|  | CV | 45.98% | 26.46% | 264.59 | 99.47% | 89.73 | 100.00% |  |  |  |  |
| F08 | mother | 54.08% | 22.47% | 128.75 | 98.80% | 0.00 | 0.00% |  |  |  |  |
|  | father | 54.18% | 22.45% | 117.40 | 98.52% | 77.56 | 100.00% |  |  |  |  |
|  | proband | 54.65% | 22.75% | 125.58 | 98.61% | 80.71 | 100.00% |  |  |  |  |
|  | plasma | 48.37% | 27.61% | 265.35 | 99.52% | 4.75 | 18.23% | 301 | 590 | 0.41% | 10.25% |
|  | CV | - | - | - | - | - | - |  |  | - |  |
| F09 | mother | 52.74% | 25.27% | 102.38 | 98.26% | 0.00 | 0.00% |  |  |  |  |
|  | father | 50.61% | 29.45% | 116.09 | 98.45% | 73.64 | 100.00% |  |  |  |  |
|  | proband | 45.86% | 24.76% | 95.38 | 97.26% | 61.80 | 100.00% |  |  |  |  |
|  | plasma | 52.43% | 28.28% | 279.38 | 99.51% | 3.15 | 10.35% | 360 | 537 | 0.40% | 15.38% |
|  | AF | 49.01% | 33.06% | 112.57 | 98.57% | 4.51 | 86.79% |  |  |  |  |
| F10 | mother | 38.78% | 36.51% | 198.08 | 99.34% | 0.00 | 0.00% |  |  |  |  |
|  | father | 37.58% | 30.05% | 162.07 | 99.23% | 107.06 | 100.00% |  |  |  |  |
|  | proband | 49.76% | 35.55% | 197.03 | 97.65% | 130.84 | 100.00% |  |  |  |  |
|  | plasma | 30.19% | 18.21% | 277.86 | 99.64% | 13.08 | 99.72% | 748 | 590 | 0.30% | 6.38% |
|  | CV | 41.24% | 37.91% | 256.23 | 98.03% | 155.85 | 100.00% |  |  |  |  |
| F11 | mother | 40.96% | 37.04% | 145.05 | 99.16% | 0.00 | 0.00% |  |  |  |  |
|  | father | 38.30% | 37.34% | 124.11 | 98.85% | 82.62 | 100.00% |  |  |  |  |
|  | proband | 41.87% | 36.14% | 122.62 | 98.81% | 80.96 | 100.00% |  |  |  |  |
|  | plasma | 29.73% | 18.13% | 240.37 | 99.66% | 17.51 | 97.15% | 782 | 855 | 0.21% | 10.93% |
|  | CV | 39.26% | 30.58% | 238.96 | 99.72% | 129.03 | 100.00% |  |  |  |  |
| F12 | mother | 51.02% | 34.25% | 135.55 | 99.09% | 0.00 | 0.00% |  |  |  |  |
|  | father | 51.06% | 36.94% | 151.68 | 99.25% | 96.51 | 100.00% |  |  |  |  |
|  | proband | 49.37% | 33.66% | 144.49 | 98.75% | 95.01 | 100.00% |  |  |  |  |
|  | plasma | 50.80% | 30.02% | 184.06 | 99.36% | 2.25 | 8.06% | 722 | 566 | 0.24% | 10.13% |
|  | CV | 36.17% | 22.14% | 227.69 | 99.59% | 2.20 | 16.28% |  |  |  |  |
| F13 | mother | 56.53% | 36.69% | 148.94 | 99.18% | 0.00 | 0.00% |  |  |  |  |
|  | father | 53.81% | 35.57% | 139.62 | 99.12% | 90.04 | 100.00% |  |  |  |  |
|  | proband | 46.23% | 34.48% | 132.11 | 99.05% | 86.90 | 100.00% |  |  |  |  |
|  | plasma | 39.51% | 14.94% | 222.78 | 99.45% | 11.32 | 97.75% | 613 | 503 | 1.31% | 3.61% |
|  | CV | 55.77% | 38.95% | 232.49 | 99.65% | 144.01 | 100.00% |  |  |  |  |
| F14 | mother | 57.70% | 30.68% | 177.84 | 99.26% | 0.00 | 0.00% |  |  |  |  |
|  | father | 52.66% | 33.75% | 181.26 | 99.37% | 117.31 | 100.00% |  |  |  |  |
|  | proband | 51.26% | 33.38% | 176.19 | 98.78% | 115.78 | 100.00% |  |  |  |  |
|  | plasma | 38.53% | 16.20% | 244.33 | 99.56% | 14.76 | 100.00% | 585 | 848 | 0.25% | 8.37% |
|  | AF | 60.35% | 35.68% | 277.83 | 99.72% | 184.41 | 100.00% |  |  |  |  |
| F15 | mother | 60.04% | 31.31% | 311.74 | 99.64% | 0.00 | 0.00% |  |  |  |  |
|  | father | 59.33% | 34.20% | 201.97 | 99.60% | 139.65 | 100.00% |  |  |  |  |
|  | proband | 59.73% | 35.01% | 252.17 | 99.14% | 171.05 | 100.00% |  |  |  |  |
|  | plasma | 41.59% | 26.02% | 224.36 | 99.58% | 14.64 | 98.94% | 764 | 659 | 0.21% | 8.23% |
|  | AF | 57.62% | 27.36% | 186.09 | 99.44% | 130.08 | 100.00% |  |  |  |  |
| F16 | mother | 61.64% | 33.26% | 228.84 | 99.52% | 0.00 | 0.00% |  |  |  |  |
|  | father | 59.79% | 26.19% | 140.40 | 99.19% | 95.96 | 100.00% |  |  |  |  |
|  | proband | 59.66% | 27.81% | 150.73 | 99.22% | 104.15 | 100.00% |  |  |  |  |
|  | plasma | 42.47% | 30.95% | 210.52 | 99.69% | 23.54 | 99.82% | 603 | 696 | 0.25% | 16.97% |
|  | AF | 58.47% | 27.85% | 195.25 | 99.55% | 131.34 | 100.00% |  |  |  |  |
| F17 | mother | 47.57% | 36.85% | 195.34 | 99.43% | 0.00 | 0.00% |  |  |  |  |
|  | father | 59.96% | 32.64% | 227.35 | 99.63% | 158.75 | 100.00% |  |  |  |  |
|  | proband | 59.96% | 31.97% | 139.82 | 99.13% | 94.01 | 100.00% |  |  |  |  |
|  | plasma | 40.39% | 33.09% | 201.42 | 99.55% | 2.66 | 12.20% | 627 | 1001 | 0.24% | 11.19% |
|  | CV | 59.18% | 28.41% | 288.07 | 99.65% | 4.73 | 55.96% |  |  |  |  |

Notes: Type 1 SNPs refers to SNPs that were homozygous with the same genotype in both parental genomes were used to calculate the plasma sequencing error rate; Type 2 SNPs refers to SNPs that were homozygous in both parents but had different types and were used to calculate the cell-free fetal DNA (cff-DNA) concentration. The CV sample in F08 family has no sequencing data due to its failed library.

In F03, F06, F07, F10, F11, F13, F14, F15 and F16, the mean depth of specific-region on Y chromosome was 17.80 (range: 11.32-26.33). The region with reads coverage at least four was over 97% (range: 97.15-100%), indicating male fetuses. In the remaining female fetuses, the mean depth of specific region on Y chromosome ranged from 2.06 to 5.87 with 4X coverage being 8.06% to 38.70%. These results were confirmed by the sequencing data of fetal genomic DNA.

We constructed fetal haplotypes for each fetus by using 348 to 977 informative SNPs phased on the target region. NIPD results revealed four normal female fetuses (F02, F05, F08 and F12), six normal male fetuses (F03, F11, F13, F14, F15 and F16), four female carriers (F01, F04, F09 and F17), and three affected male fetuses (F06, F07 and F10) (Table 3 and Figure 3). The result was in concordant with invasive diagnosis using MLPA.

**Figure 3.**
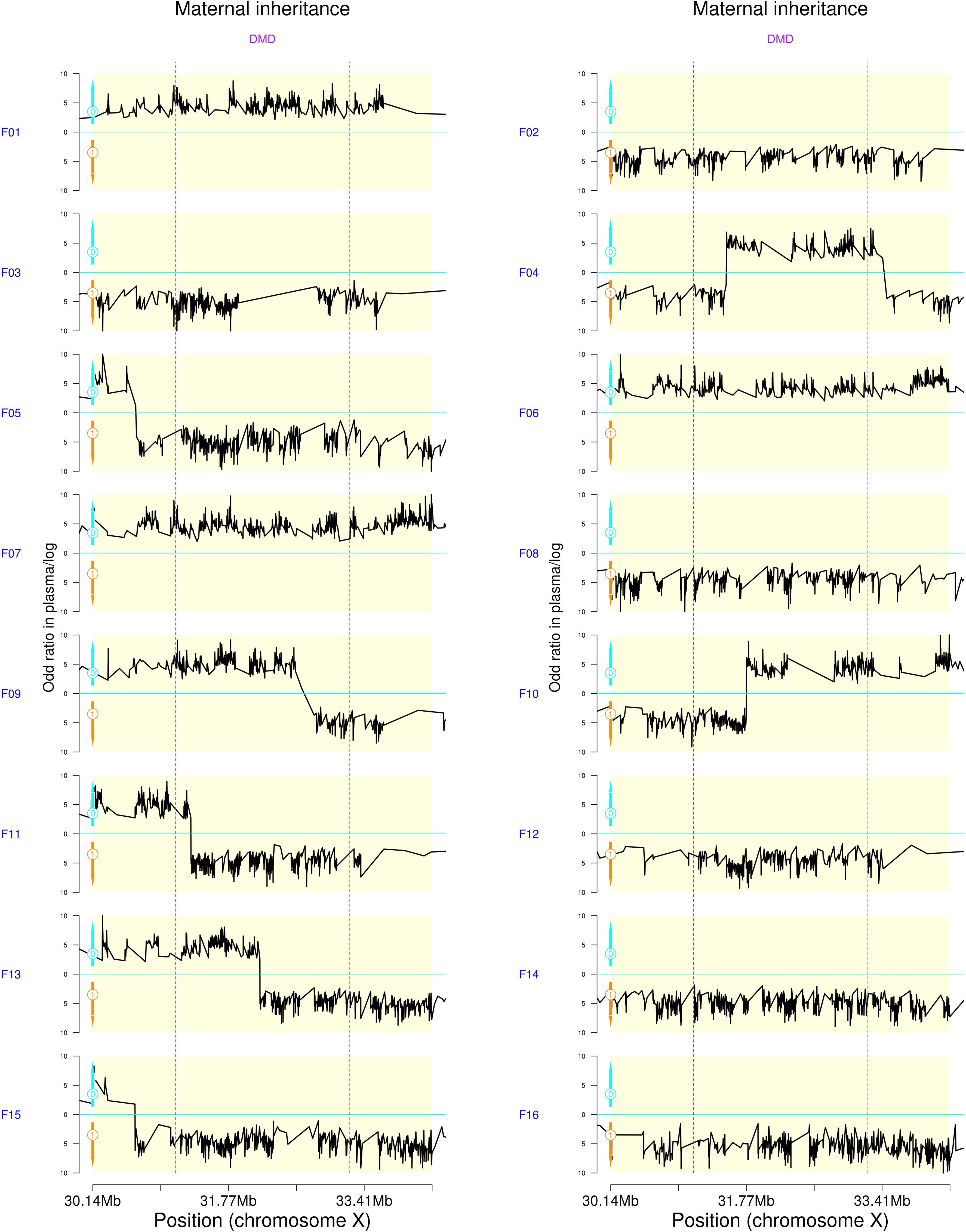
Fetal haplotype inference. X-axis represents the locus on chromosome X, Y-axis represents the logarithm of the ratios of fetal different haplotype combinations; the blue Line fetus inherited from maternal haplotypes. Lines above zero (Cyan lines) point to that the fetus inherited the pathogenic allele (Hap0), and the lines below zero point to that the fetus inherited the normal allele (Hap1).

**Table 3.** Noninvasive Prenatal Test of DMD Families

| Family | Fetal Gender | Phased SNPs<sup>a</sup> | SNPs For M0<sup>b</sup> | SNPs For M1<sup>c</sup> | Inherited Maternal | Predicted results | NIPT-based Test | MLPA diagnosis |
| --- | --- | --- | --- | --- | --- | --- | --- | --- |
| F01 | Female | 563 | 560 | 0 | M0 | Carrier | Het.<br>D.EX46-48 | Het.<br>D.EX46-48 |
| F02 | Female | 554 | 0 | 551 | M1 | Normal | Normal | Normal |
| F03 | Male | 445 | 0 | 444 | M1 | Normal | Normal | Normal |
| F04 | Female | 402 | 234 | 168 | M0 | Carrier | Het.<br>D.EX43-45 | Het.<br>D.EX43-45 |
| F05 | Female | 469 | 29 | 440 | M1 | Normal | Normal | Normal |
| F06 | Male | 442 | 441 | 0 | M0 | Affected | D.EX49-52 | D.EX49-52 |
| F07 | Male | 631 | 629 | 0 | M0 | Affected | Hem.<br>c.628G>T | Hem.<br>c.628G>T |
| F08 | Female | 585 | 0 | 582 | M1 | Normal | Normal | Normal |
| F09 | Female | 434 | 310 | 123 | M0 | Carrier | Het.<br>D.EX45-48 | Het.<br>D.EX45-48 |
| F10 | Male | 610 | 350 | 253 | M0 | Affected | D.EX8-37 | D.EX8-37 |
| F11 | Male | 905 | 231 | 654 | M1 | Normal | Normal | Normal |
| F12 | Female | 424 | 0 | 423 | M1 | Normal | Normal | Normal |
| F13 | Male | 783 | 275 | 505 | M1 | Normal | Normal | Normal |
| F14 | Male | 977 | 0 | 962 | M1 | Normal | Normal | Normal |
| F15 | Male | 599 | 13 | 575 | M1 | Normal | Normal | Normal |
| F16 | Male | 529 | 0 | 527 | M1 | Normal | Normal | Normal |
| F17 | Female | 348 | 339 | 0 | M0 | Carrier | Het.<br>D.EX14-15 | Het.<br>D.EX14-15 |

Notes: M0, fetal-inherited maternal mutant haplotype; M1, fetal-inherited maternal wild-type haplotype;
SNPs<sup>a</sup> represents SNPs that were used to predict maternal haplotypes with a trio strategy.
SNPs<sup>b</sup> represents number of SNPs supported for M0;
SNPs<sup>c</sup> represents number of SNPs supported for M1.

To further evaluate the inferred fetal haplotype, we performed target capture sequencing in CV or AF samples (failed in CV in F08). Fetal haplotypes constructed using the parental-fetus genomic DNA sequencing data were same as the corresponding fetal haplotypes inferred through plasma DNA sequencing. There were no false positive/negative results.

## Discussion

In this study, all 17 fetuses were accurately diagnosed as normal, carrier or affected using haplotype-based NIPD. This study and previous works of NIPD for DMD showed a high degree of accuracy compared with the invasive diagnosis results, as shown in Table 4[11-14]. We also reviewed NIPD for other monogenic disorders[15]. The accuracy of NIPD for SGDs was near 100%[6, 7, 14, 16-21] (Table 5).

**Table 4.** Studies reporting the accuracy of non-invasive prenatal testing of DMD

| Author |  | Country | Research Design | Blind method | Objects | Gold standard of Diagnosis | Number of case | of A c |
| --- | --- | --- | --- | --- | --- | --- | --- | --- |
| Yan Xu |  | China | Prospective Study | Yes | DMD/BMD families | Sanger sequencing/quantitative real-time PCR | 8 | 1 |
| Michel Parks | 10 | UK | Prospective Study | Yes | Aneuploidy families/DMD families | Amplification analysis (MLPA) | 9 | 1 |
| Seong-Keun Yoo | 15 | Korea | Prospective Study | Yes | DMD/BMD families | -- | 4 | 1 |
| Current Study |  | China | Prospective Study | Yes | DMD/BMD families | Amplification analysis (MLPA) | 17 | 1 |

Only first author is given for each study.

**Table 5.**
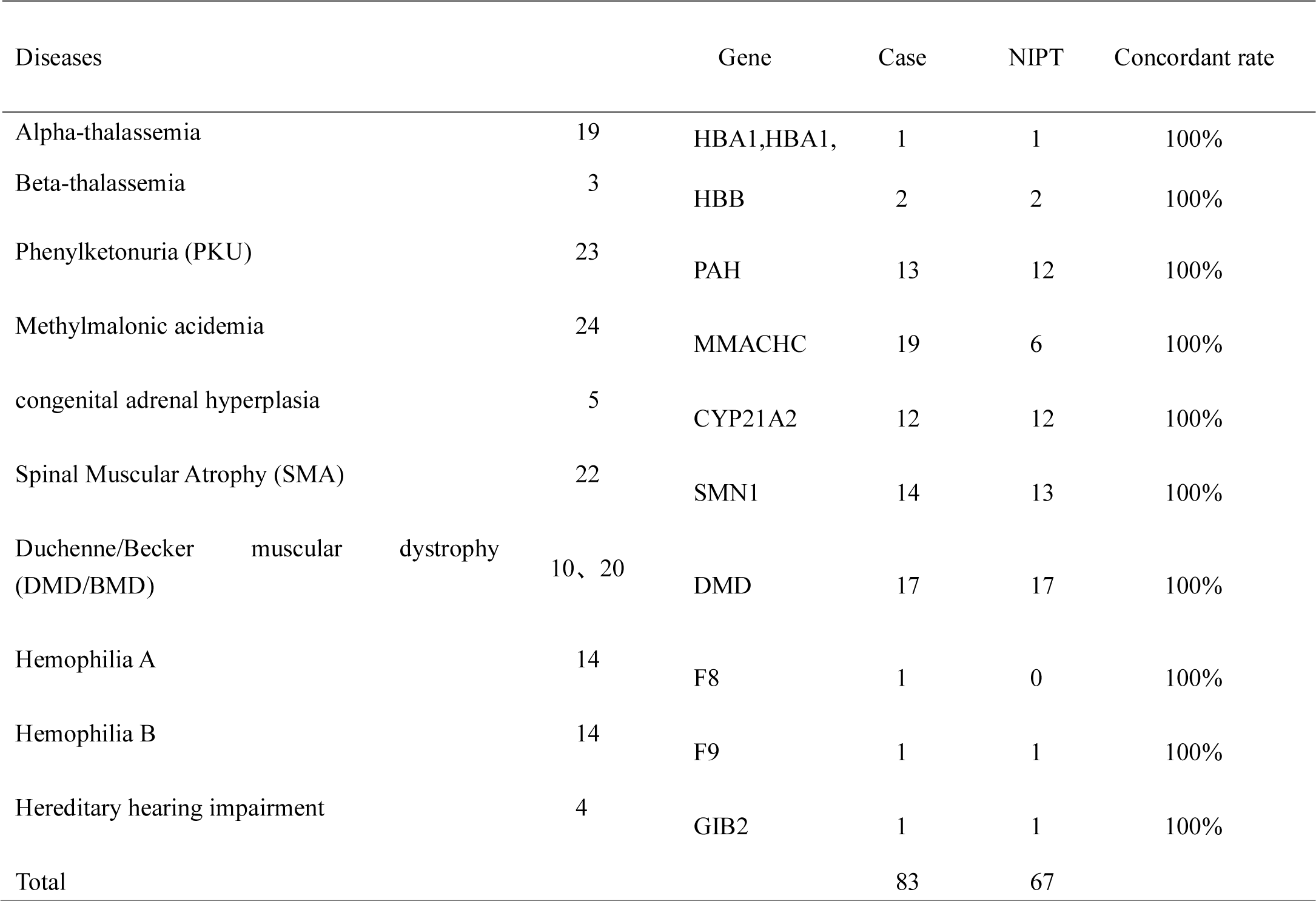
The accuracy of NIPT for other single-gene diseases

Although technically possible, the high cost of NIPD may be a major factor limiting its clinical application. The cost is about 600-900 dollars for one sample based on an independent experimental operation. The cost is mainly composed of high depth sequence, the commercial capture probe and bespoke design service for specific gene and mutation uncovered in existing probe. The small sample size in one turn round of NIPD would also increase the cost. In our work, several attempts have been made in order to reduce the cost. First, the capture region of the customized probe was narrowed to 657.29 kb including the coding region and flanking region in *DMD* gene. Second, 100x sequence depth was determined to be cut off to balance the relationships between sequence cost and accuracy. Third, explore the maximization of samples on the same probe capture reaction. In our study, the best experimental scheme was to recruit samples of five families, each of which included maternal cff-DNA, the parental and proband’s genomic DNA (gDNA). The capture probe could be expanded to more monogenic diseases with higher incidence to reduce the bespoke design cost. With the development of technology[22], the cost of sequencing will continue to decrease.

The sequencing depth, fetal fraction and number of informative SNPs are key factors to ensure the accuracy and reliability of NIPD in clinical service. Our study innovatively elaborated quality control using the computer simulation. When the fetal fraction is below 5%, the number of corresponding SNPs required for fetal haplotype constructing is at least 40 to reach the accuracy of 99%. When the plasma sequence depth is 200x, the accuracy of inferred fetal haplotype is close to 100% (Supplemental Fig S1). Fetal fractions of other families were all above 5%, the informative SNPs were all above 200 and the accuracy of NIPD was 100%. If the informative SNPs were enough to predict the fetal haplotype, we could reduce the plasma sequence depth to cut the cost. So, we evaluated the effect of sequencing depth on inferring fetal haplotypes (Supplemental Fig S2). If the sequence depth of plasma was reduced to 100X, 20 or more SNPs were used to infer fetal maternal haplotypes to achieve an accuracy of 99%. If the sequence depth of plasma was reduced to 60x, at least 35 SNPs was needed to predict the fetal inherited maternal haplotype to ensure an accuracy of 99%. The accuracy depended on the number of informative SNPs affected by sequence depth of plasma and fetal DNA fraction. Therefore, the cut-off must be determined by considering the interaction of three factors. According to the computer simulation experiments, to ensure the detection accuracy with 99%, the cutoff value of the number of informative SNPs was 20, the dedup depth of plasma was prescribed as 100x and the fetal DNA fraction was defined to be 5%.

Due to the recombination frequency of the entire dystrophin gene can be as high as 12%[23], it is crucial to infer the fetal inherited allele for recombination accurately. Currently, the position of the recombination event was confirmed by comparing the haplotypes of the proband and the CVS, or by bi-direcional (i.e. 5’to 3’and 3’to 5’) evaluation of recombination sites in RHDO analysis[16]. We precisely assessed the position of the combination site to support a correct diagnosis, demonstrating the robustness of the haplotype-based approach. Nevertheless, further improvements are necessary. For instance, coverage of SNPs across various autosomes should be, increased to further improve the accuracy. Establishing referral laboratories for DMD with increased multiplexing capacity of multi-personnel in each sequencing run will help to reduce the cost and shorten the turn-around-time[16].

Recently, linked-reads sequencing technology based on microfluidics has become available, allowing direct haplotype phasing of the target region and NIPD were successfully achieved[13]. This approach does not rely on the availability of DNA from affected proband and should be accessible for more couples. However, the cost of linked-reads technology was three times as our method due to its expensive library kits and equipment. The 10x Genomics technology was more labour intensive and not asreadily scalable as our method. Its application in the high-risk pregnancy for DMD needs further study.

## Acknowledgments

We thank all participants in this study for their collaborations. This work was supported by the National Natural Science Foundation of China (NSFC) (No. 81671470), Guangzhou Science and Technology Program (No. 2014 Y2-00551, No. 201504282321393, No. 201604020078, No.201604020091), Guangdong Science and Technology Program (No. 2013B022000005, No. 2016A030313610), and Major Technical Innovation Project of Hubei Province (No. 2017ACA097).

