## Supplementary figures and images for "Haplotype-Based Noninvasive Prenatal Diagnosis for Duchenne Muscular Dystrophy: A pilot study in South China"

### Supplemental Figure 1

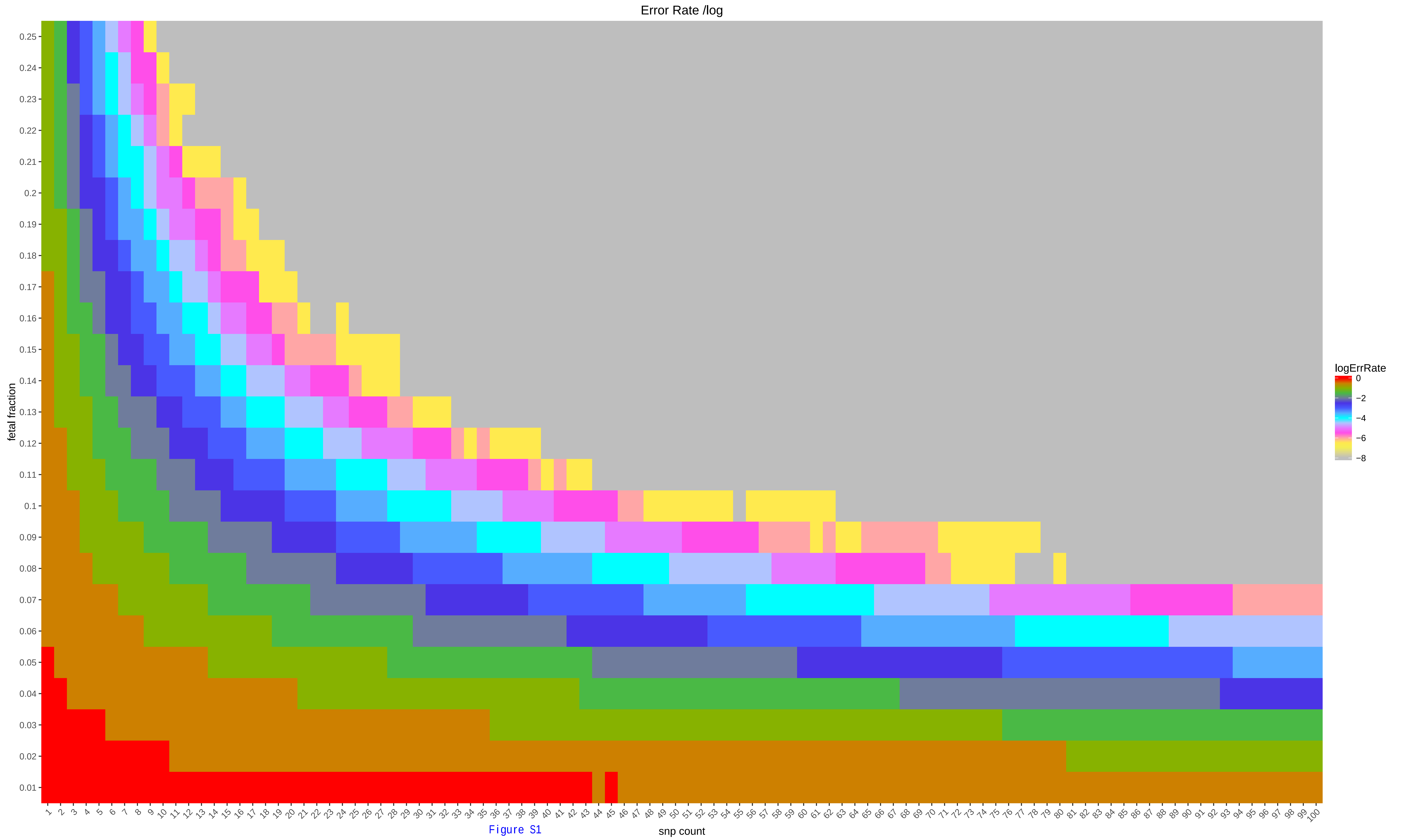

Figure S1

### Supplemental Figure 2

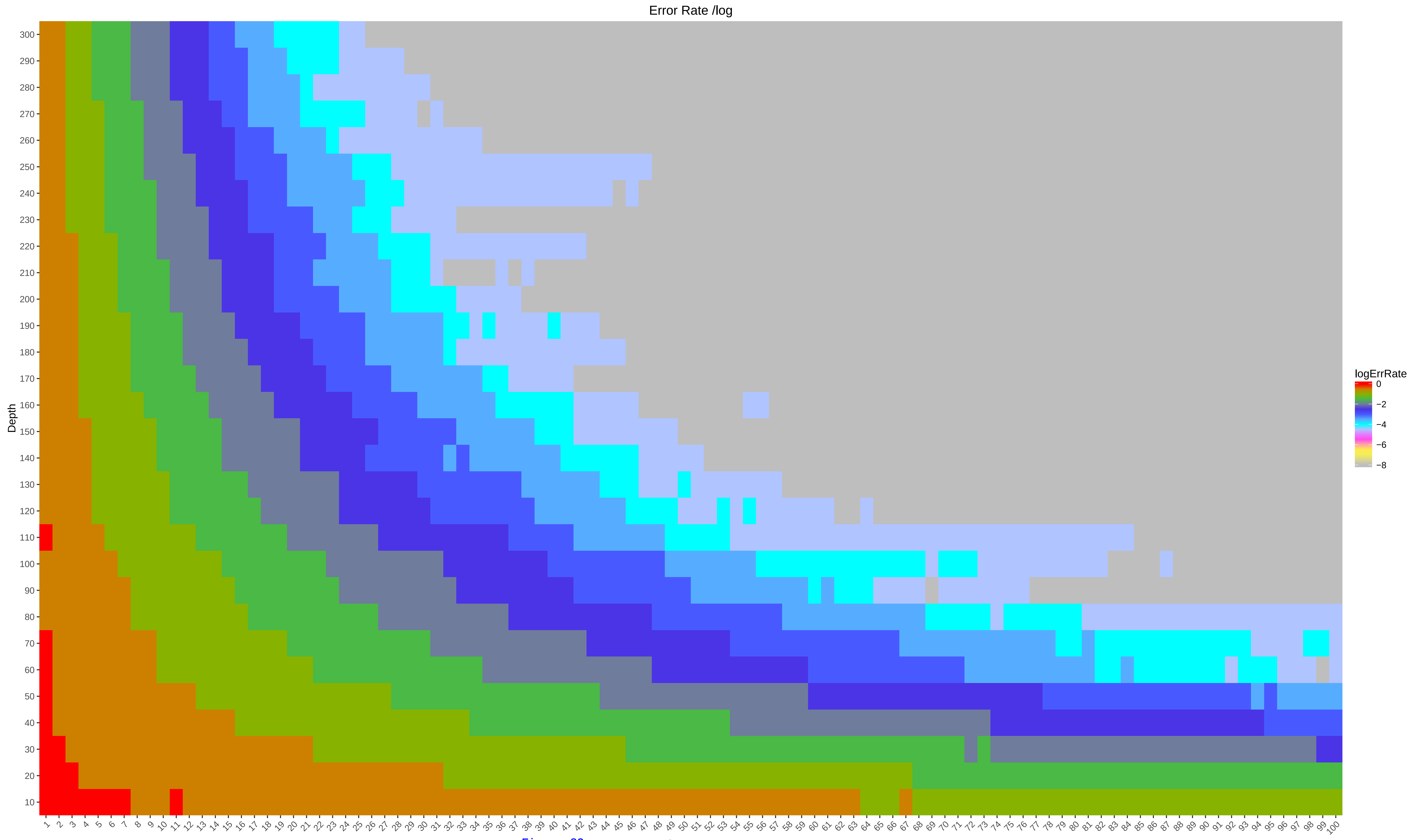

Figure S2
